# The Uniqueness of Human Vulnerability to Brain Aging in Great Ape Evolution

**DOI:** 10.1101/2022.09.27.509685

**Authors:** Sam Vickery, Kaustubh R. Patil, Robert Dahnke, William D. Hopkins, Chet C. Sherwood, Svenja Caspers, Simon B. Eickhoff, Felix Hoffstaedter

## Abstract

Aging is associated with robust decline of the brain’s gray matter. This spatially specific, morphological change in humans has recently been found in chimpanzees. Direct comparison of age-related brain deterioration between these great ape species can provide a unique evolutionary perspective on human brain aging. Here, we present a data-driven, cross-species comparative framework to explore the relationship between gray matter atrophy with age and cross-species cerebral expansion in chimpanzees and humans. In humans, we found a positive relationship between cerebral aging and cortical expansion, whereas, in chimpanzees no such relationship was found. The greater aging and expansion effects in higher-order cognitive regions like the orbito-frontal cortex were observed to be unique to humans. This resembles the last in, first out hypothesis for neurodevelopment and may represent a biological cost for recent evolutionary developments of human faculties.

## Introduction

With age, pronounced alterations occur in morphology and organization of the human brain with a distinct spatial pattern resulting in part from cellular atrophy in later life^1,2^. This aging process may be further accelerated by age-mediated disorders such as Alzheimer’s disease, Parkinson’s disease, and other neurodegenerative conditions^3^. Furthering our understanding about specific neurobiological influences on spatial patterns of brain aging, may provide insight into the brain changes in healthy aging and possible diagnostic markers for clinical outcomes. Historically, comparative neuroscience has been an effective catalyst for important discoveries regarding principles of anatomy and functional specializations of the human brain^4^. With open and collaborative endeavors such as the National Chimpanzee Brain Resource (NCBR) and the PRIMatE Date Exchange (PRIME-DE)^5^, along with improved methodologies and imaging technology, large scale comparative neuroanatomy has become able to answer new translational questions^6^.

Morphological gray matter (GM) changes during aging have recently been shown to be present in one of humans’ closest extant primate relative, chimpanzees (*Pan troglodytes*)^7,8^, where age-related changes are similar but at a lower magnitude compared to humans^8^. For example, age-related volumetric reduction of overall hippocampus and frontal cortex size are not evident in chimpanzees, but occurs in humans that could be related to humans comparatively extended lifespan^9^. Cognitive decline is also present in chimpanzees, but appears not as prominent as in humans^10^. In this context, understanding GM alterations during brain aging in great ape evolution (e.g. which includes humans and chimpanzees, as well as bonobos, gorillas, and orangutans) may aid in understanding the spatial distribution of morphological changes due to healthy aging and disease.

The comparison of neuroanatomy and brain functions across primate species is commonly informed by analyzing homologous brain regions^6,11–13^. Classically, these regional homologies are defined by manually delineating brain partitions, based on macroanatomy, gene expression, connectivity, and/or cytoarchitectonic features. This approach rests on the assumption that similar anatomical features result in a common functional organization across species and thereby enable an informative and meaningful comparison between them. However, such homologies can be contentious, for example the delineation of the prefrontal cortex (PFC) in primates^11^ and therefore can be influenced by methodological biases^14^. The homologous-centric approach has proven to be effective and informative. Utilizing a data-driven approach can supplement these techniques while still capturing important cross-species differences and incorporating species-specific features in a datacentric manner^15^.

Chimpanzees offer an ideal referential model to investigate evolutionary changes within the human lineage as they share a last common ancestor with humans approximately 6 - 8 million years ago^16^. Accordingly, chimpanzees and humans have substantial genomic similarities^17^ as well as neuroanatomical features in common^18–20^. Furthermore, new evidence suggests that menopause occurs at a similar age in humans and chimpanzees, with demographic and hormonal data indicating that reproductive cessation in both species is caused by a common physiological factor^21^. Consequently, chimpanzees represent a unique possibility to infer distinctive evolutionary adaptations of the human brain by analyzing commonalities and recent divergences, although current neurobiology also reflects species-specific adaptations. Previous studies have shown that multimodal association cortices in humans are disproportionately larger than in non-human primates^11,22,23^. The higher expansion of certain brain areas through human evolution likely relate to human specific cognitive functions. Specifically, the greatly expanded human PFC^11^ can be associated with self-control and executive functioning^24^ and the larger precuneus^22^ with visuospatial processing^25^. Furthermore, the vulnerability of frontal cortical areas to aging processes is hypothesized to be related to their late maturation^26^. This refers to the “last in, first out hypothesis” and interestingly similarities have been shown between cortical development and cross-species expansion^27^.

In this study, we directly compare age-mediated GM changes in chimpanzees and humans, which represent two species in the Hominidae family (i.e., great apes) and explore their relationship with cross-species cerebral expansion. For inter-species comparison, we developed a multivariate data-driven comparative framework that applies voxel-wise clustering based on GM variability within each species independently. The optimal low dimensional representation of brain morphology for each species is then compared in a cross-species investigation of aging and brain expansion. Comparative data for calculating cross-species expansion was provided via proximate phylogenetic ancestors. Accordingly, humans were compared to chimpanzees, while chimpanzees were compared to olive baboons (*Papio anubis*) and rhesus macaques (*Macaca mulatta*), two commonly researched cercopithecoid monkey species. Thus, we test whether the relationship between aging and cerebral expansion is unique to humans or instead might be a feature shared between humans and chimpanzees possibly originating at the divergence of the great ape lineages from other primates.

In summary, we present a novel approach for data-driven cross-species comparison of structural brain organization and demonstrate its utility by analyzing the relationship between cerebral aging and cross-species expansion in humans and chimpanzees. Our data-driven approach uses both species-specific information in addition to cross-species similarity to create an anatomically interpretable low-dimensional brain parcellation. We show that the resulting parcellation aligns with known macroanatomical structures in both humans and chimpanzees. Applying this comparative framework, we jointly analyze spatial pattern in brain aging and cerebral expansion of the two great ape species. Finally, we present evidence for a relationship between local age-mediated GM changes and recent cortical expansion in humans that is not present in chimpanzees.

## Results

Our cross-species comparative approach was based on structural magnetic resonance imaging (MRI) scans from 189 chimpanzees and 480 human brains (Fig. 1B). Orthogonal projective nonnegative matrix factorization (OPNMF)^28,29^ was applied to normalized GM maps within each species independently. The orthogonality and non-negativity constraints of OPNMF results in a spatially continuous, parts-based representation of the input data based on regional covariance of brain structure within each species^30^. OPNMF has been extensively used with human neuroimaging data yielding anatomically meaningful correspondence of clustering solutions^28,31–34^.

**Figure 1.**
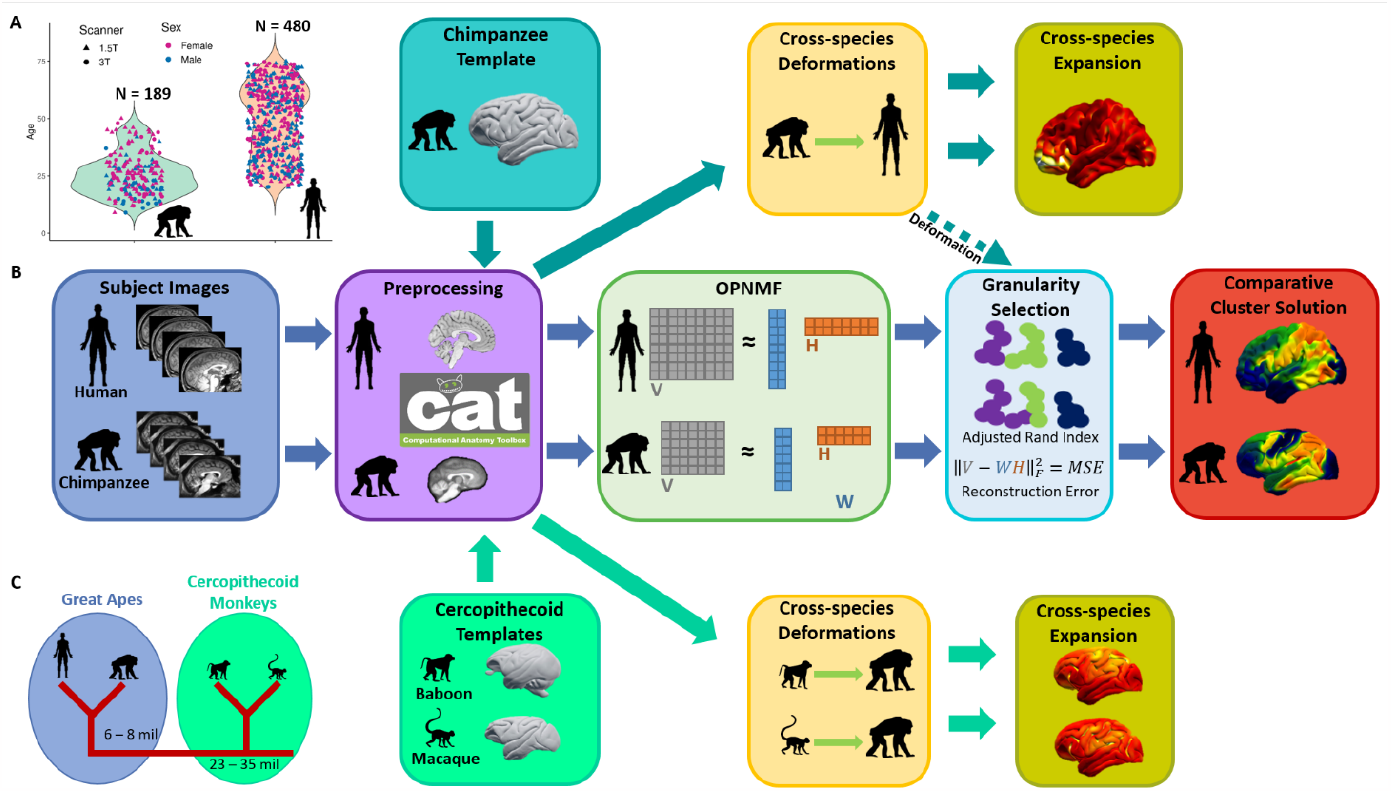
Sample, Workflow and Phylogeny. A – Age, sex, and scanner field strength distribution of the chimpanzee (N=189) and human (N=480) samples. B – Workflow outlining our comparative approach by utilizing OPNMF and creating cross-species expansion maps. C – Diagram showing the phylogenetic relationship of humans to the other three primate species investigated in this study.

The comparative framework utilizing OPNMF as well as the creation of the cross-species expansion maps is outlined in Figure 1A. The approach begins with separately segmenting and normalizing the individual chimpanzee and human images utilizing species-specific templates in almost identical CAT12 pipelines^8,35^. The processed GM maps for each species are parcellated independently using OPNMF over a range of granularities (2-40) and bootstrapped over the whole sample with replacement to ensure stability of the solutions. Mean reconstruction error (MRE) over bootstraps is used to select a range of clustering solutions with optimal numbers of parcels for cross-species comparison. For direct cross-species comparison, the JunaChimp average chimpanzee T1-weighted (T1w) template^8^ is submitted to the human preprocessing pipeline to create a representative chimpanzee to human deformation field. The JunaChimp^8^ to MNI152 (ICBM 2009c Nonlinear Asymmetric^35^) deformation map is used to non-linearly register the chimpanzee OPNMF solutions to the human template space for the analysis of parcel or factor similarity using the adjusted rand index (ARI). These cross-species parcel similarity of multivariate GM morphology are used for the selection of optimal parcellation granularity, together with species-specific OPNMF MRE. To create cross-species expansion maps for chimpanzees, average population templates from olive baboon^36^ and rhesus macaque^37–39^ were processed with the chimpanzee^8^ pipeline.

### Comparative Brain Parcellation

OPNMF clusters the volumetric GM maps and yields parcels which contain voxels that co-vary with on another across the sample. This unsupervised clustering technique behaves similar to other ^40^ like independent component analysis and requires an a priori decision on the number of cluster/parcels to represent the original data^28^. The decision for the most appropriate OPNMF solution was determined via assessing cross-species spatial similarity and the development of OPNMF reconstruction accuracy at different granularities (Fig. 2A). Chimpanzee parcellations were transformed to the human template space using the chimpanzee to human deformation map for assessing cross-species parcel similarity using ARI. Quality control of the chimpanzee to human deformation map was conducted by visually inspecting the overall fit of the Davi130 chimpanzee macroanatomical labels^8^ projected to human template space (Supplementary Figure 1). The OPNMF solution with highest ARI for within species parcellations represents common cross-species organizational patterns of GM covariance^30^. The MRE indicates how accurately the input data (GM maps) can be represented by the OPNMF factorization. By increasing the number of clusters, more variance in GM input data is modeled and MRE naturally decreases, while this association is non-linear and sample specific (Fig. 2A). A plateau of the MRE decrease with increasing OPNMF granularity only marginally improves the solution’s representation of GM data. Consequently, the beginning of a plateau hints at a good tradeoff between the solutions reconstruction accuracy and the complexity of the cluster solution. Finally, to ensure the robustness of the MRE development curve, 100 bootstraps were computed for each OPNMF granularity in both species.

**Figure 2.**
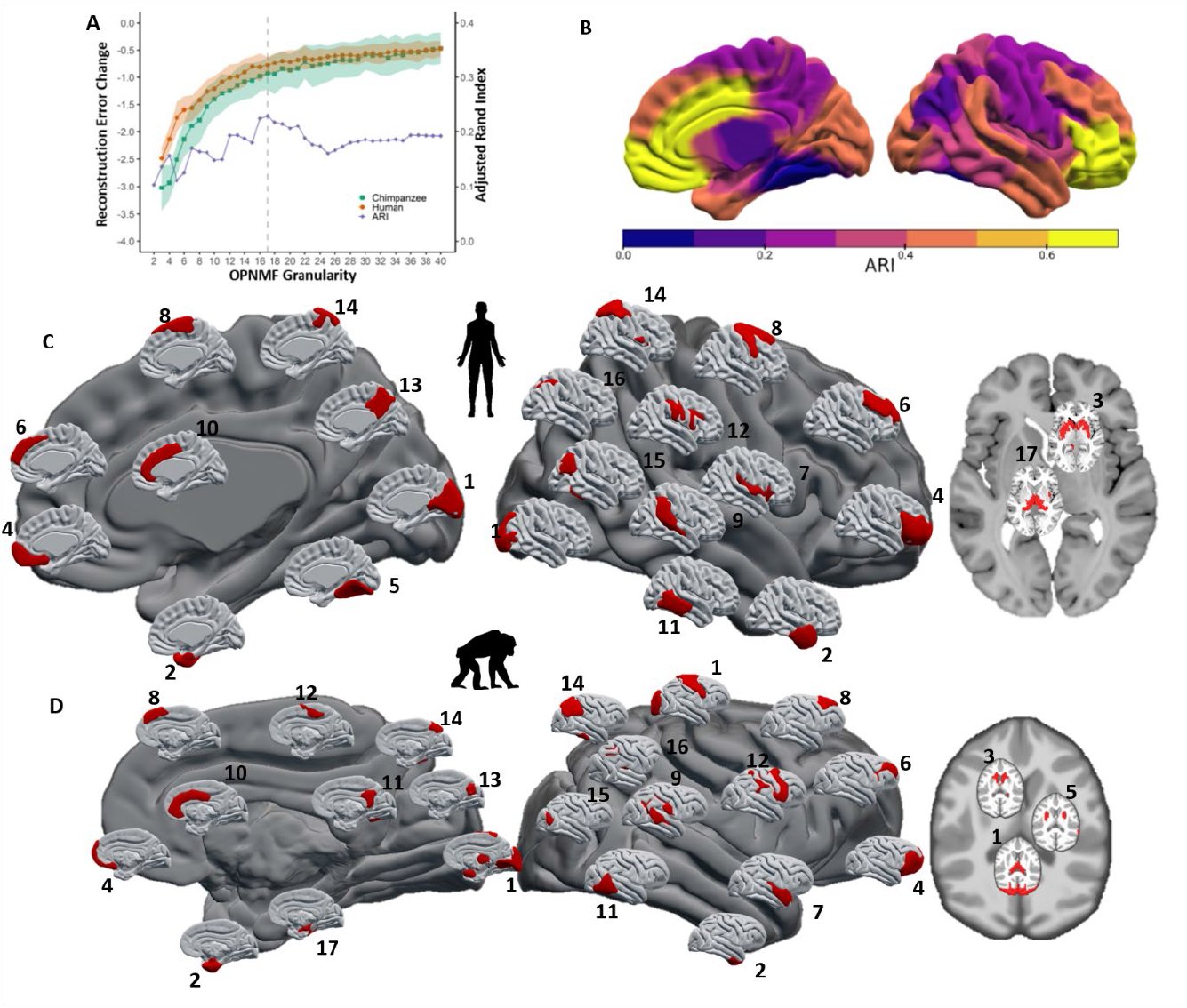
17-cluster OPNMF solution for cross-species comparison. A – OPNMF granularity selection utilizing ARI to assess cross-species similarity and relevant change in reconstruction error (RE) over a granularity range of 2 – 40 clusters and bootstrapped (k=100) to ensure stability. The one standard deviation from the change in MRE over 100 bootstraps is represented as shadow, the gray dashed line represents the selected number of 17 clusters. B –Cross-species single parcel ARI in human template space. C – Human selected 17-cluster OPNMF solution with macroanatomical labels: 1 – occipital lobe, 2 – temporal pole, 3 – putamen, caudate nucleus, amygdala, and hippocampus, 4 – prefrontal and orbito-frontal cortex, 5 – lingual and fusiform gyrus, 6 – superior and middle frontal gyrus, 7 – insula, 8 – pre-central gyrus and pre-motor area, 9 – temporal parietal junction, 10 – anterior and middle cingulate cortex, 11 – posterior middle and inferior temporal gyri, 12 – supramarginal gyrus, inferior post-central gyrus, and inferior pre-central sulcus, 13 – precuneus, 14 – superior parietal lobe, 15 – angular and fusiform gyrus, 16 – superior parietal sulcus and parahippocampal cortex, 17 – thalamus. D - Selected 17-cluster OPNMF solution for chimpanzees with macroanatomical labels: 1 – occipital lobe, primary motor cortex, and thalamus, 2 – temporal pole, 3 – caudate nucleus, 4 – prefrontal and orbito-frontal cortex, 5 – putamen, 6 – middle frontal gyrus, 7 – superior temporal gyrus and anterior insula, 8 – posterior superior frontal gyrus, 9 – temporal parietal junction and supramarginal gyrus, 10 – anterior and middle cingulate cortex, 11 – posterior cingulate, precuneus, and peristriate cortex, 12 – supplementary and pre-motor areas, 13 – cuneus and medial occipital-parietal sulcus, 14 – superior and inferior parietal lobe and inferior temporal gyrus, 15 – lateral parietal-occipital sulcus, 16 – superior parietal sulcus and posterior insula, 17 – amygdala and hippocampus.

The highest spatial similarity of parcellations between species was found for the 17-cluster solution with a mean ARI of 0.23 (Fig. 2A) including several parcels with ARI > 0.4 (Fig. 2B). The MRE development curve did not show a clear indication of a plateau for both species, yet a gradual plateau is present in the range of 15 – 21 clusters in chimpanzees and 14 – 20 in humans. Therefore, the 17-cluster solution met our criteria for both chimpanzees (Fig. 3A) and humans (Fig. 3B) for a data-driven cross-species comparative investigation.

**Figure 3.**
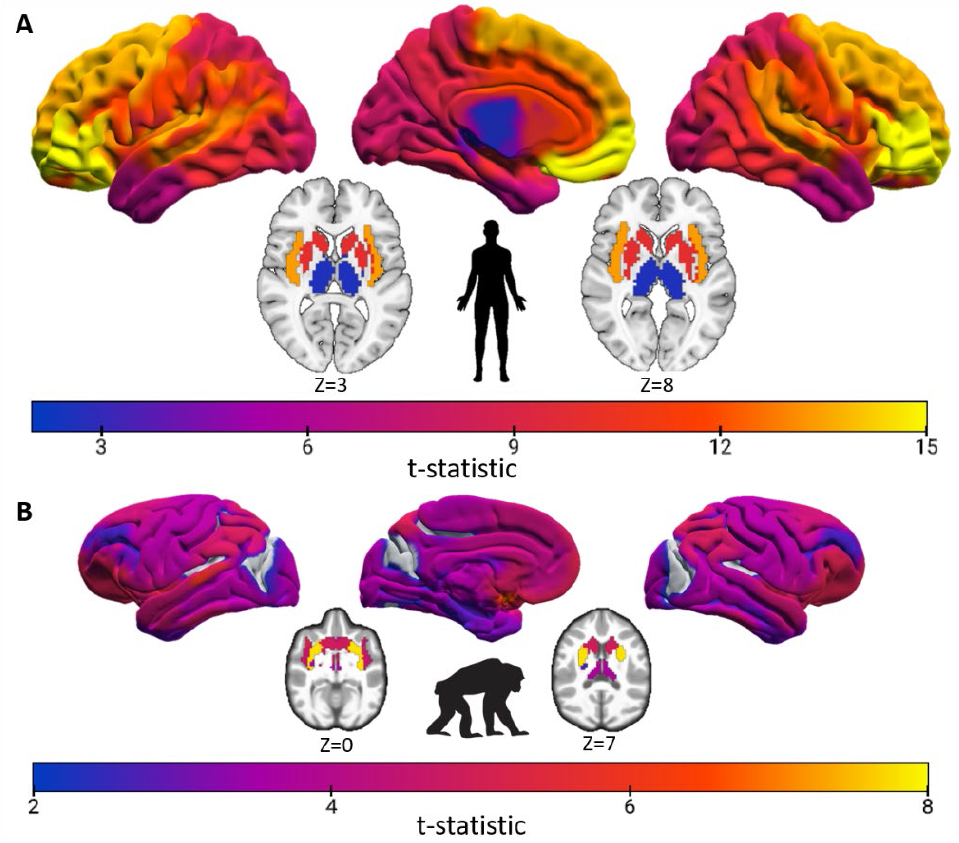
Age related GM decline in OPNMF 17-cluster solutions. Age-mediated GM changes presented as cluster-wise regression model absolute t-statistics FWE corrected p≤0.05 in A - chimpanzees and B - humans.

The 17-cluster OPNMF solutions in both chimpanzees (Fig. 2D) and humans (Fig. 2C) represents a data-driven parcellation of both species cerebral GM. These regions were established through a hard parcellation by assigning each voxel to the parcel with the highest weight enforced by the orthogonality constraint of OPNMF. Some resulting parcels closely align with known macroanatomical regions in both species like the orbito-frontal cortex, middle frontal gyrus, anterior and middle cingulate cortex, and the temporal pole (Fig. 2C&D). In contrast, for humans separate parcels delineated the insula, superior parietal lobule, precuneus, occipital lobe, and thalamus (Fig. 2D), while in chimpanzees the pre-motor cortex, hippocampus, putamen, and caudate nucleus were differentiated (Fig. 2C). Overall, orbito-frontal and the cingulate cortices showed the highest crossspecies similarity with an ARI of 0.66 and 0.64 respectively (Fig. 2B). A marked difference between species can be seen in the parcellation of sensory-motor cortices. In chimpanzees, two parcels represented major sensory-motor structures, one for the occipital lobe, pre- and post-central gyrus, and thalamus and another for the pre-motor cortex. In humans, separate parcels represented thalamus and occipital lobe as well as motor and pre-motor cortices. Specifically in humans, multimodal parietal regions like the precuneus, superior parietal lobule, angular gyrus, and temporal-parietal junction were parcellated into different clusters, while in chimpanzees the basal ganglia are more differentiated.

### Brain Aging

Age-mediated GM decline in chimpanzees and humans was assessed for the OPNMF 17-cluster solution. The average GM volume of each parcel was used as the dependent variable in a multiple linear regression model with age, sex, total intracranial volume (TIV), and scanner field strength as independent variables. To improve comparability, the human sample age range was matched to the chimpanzees by accounting for the interspecies differences in brain aging. The comparative aging difference of human years approximated to 1.15 years in chimpanzees was used based on a comprehensive study using a combination of anatomic, genetic, and behavioral data^41^. Accordingly, as the oldest chimpanzees were 50 years old, humans over 58 years old were removed to include 304 subjects (150 females; mean age = 39.0±11.0) for the age regression model. Of note, this represents a middle-aged human sample, including minimal morphological changes due to age-related neurodegenerative or pre-clinical conditions such as mild cognitive impairment. In both species, significant age-mediated GM decline was found in nearly all parcels following correction for multiple comparisons across parcels at p≤0.05^42^. Humans showed age-related GM decline across all parcels, largest in frontal and prefrontal cortices. Chimpanzees displayed significant age-mediated GM decline in all but three parcels, which represent the peristriate cortex, posterior insula, cuneus, and superior parietal sulcus. Both species showed relatively low age related changes in occipital and motor areas. The largest age-related GM decline in chimpanzees was found in the striatum, in particular the caudate nucleus. Of note, comparable spatial distribution of age-related GM decline was found when employing the same macroanatomical Davi130 parcellation^8^ with 7.6-fold higher granularity in both chimpanzees and humans (Supplementary Figure 2).

### Cross-species Brain Expansion

Utilizing the 17-cluster solution, we compared cross-species brain expansion based on population representative T1w templates from humans^43^, chimpanzees^8^, olive baboons^36^ and rhesus macaques^37–39^. Template images provide artificially high tissue contrasts in addition to representing an average brain of a particular species aiding generalizability. To provide estimations for cerebral expansion, we calculated cross-species non-linear coregistration of the brain from chimpanzee to human, baboon to chimpanzee, and macaque to chimpanzee. We present the estimated cross-species expansion maps as relative expansion, where values of one represents a local volumetric expansion comparable to the overall difference in brain size between species. Therefore, values greater than one represent local brain areas that have expanded more than the overall difference in brain size between species and less than one show lower expansion than brain size difference.

The largest human expansion was found in the orbito-frontal cortex, which additionally showed the greatest age-mediated GM decline (Fig. 3B) and also the highest cross-species parcel similarity (Fig. 2B). Further large cortical expansion was found in other multimodal association areas such as the middle and medial frontal cortex, superior parietal, precuneus, insula, and cingulate cortex and low expansion was located in the temporal pole as well as occipital, motor, and subcortical areas (Fig. 4A). These latter regions also contained lower expansion in the baboon (Fig. 4B) and macaque (Fig. 4C) to chimpanzee expansion maps, although, the cercopithecoid monkeys to chimpanzee presented much lower expansion as compared to the chimpanzee to human map. Additionally, the precuneus showed high expansion in the human (Fig. 4A) and relatively low in the chimpanzee from both cercopitheciod monkeys. The general pattern of expansion in both cercopithecoid monkeys to chimpanzee is similar, with large expansions of frontal, parietal, and cingulate cortices. In baboon to chimpanzee, the largest expansion occurred in the superior frontal gyrus/pre-motor area (Fig. 4B) while in macaque to chimpanzee, the superior parietal sulcus and posterior insula (Fig. 4C) featured the largest expansion. Furthermore, macaque to chimpanzee showed comparably more expansion in the occipital-parietal junction and lower expansion in the motor/pre-motor area, occipital cortex, and basal ganglia compared with baboon to chimpanzee expansion. To summarize, chimpanzee to human as well as cercopithecoid monkeys to chimpanzee expansion maps show relatively high expansion in frontal and parietal cortical regions. The human features the greatest expansion in prefrontal areas while in chimpanzees the largest expansion was seen in pre-motor/frontal and lateral parietal regions.

**Figure 4.**
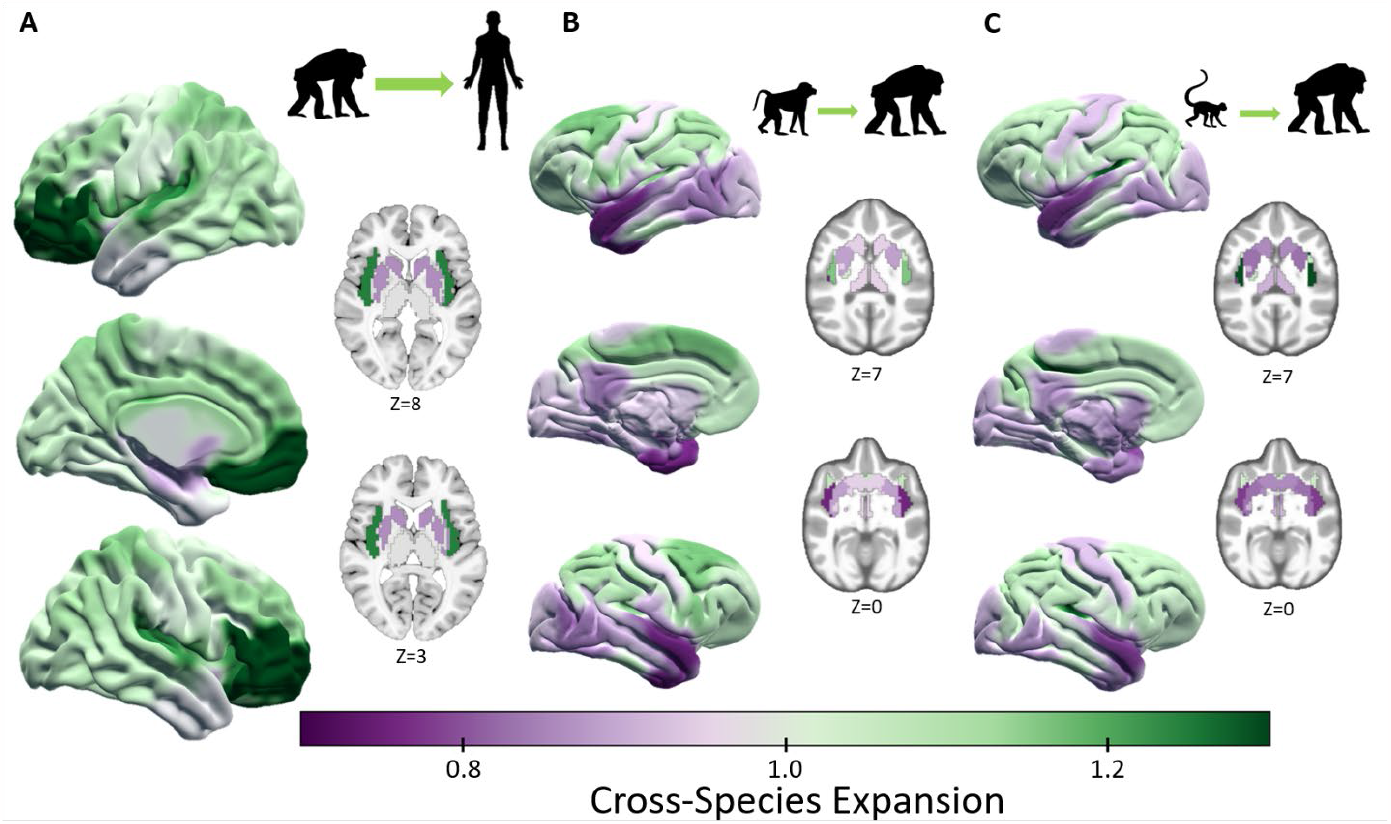
Species specific OPNMF 17-clusters of cerebral expansion. Average relative expansion for each OPNMF 17-cluster is shown for A - human expansion from chimpanzee B - chimpanzee from baboon, & C - chimpanzee from macaque expansion.

### Brain Aging and Cross-species Expansion

We investigated the relationship between cross-species expansion and age-related GM changes in chimpanzees and humans. For humans, brain aging was compared with cortical expansion from chimpanzee to human while for chimpanzees, aging was compared with expansion relative to both baboon and macaque. A strong positive correlation was found between cerebral expansion and age-mediated GM decline in humans (Fig. 5A), following permutation testing at p≤0.05 (r = 0.52; p = 0.01). This relationship is particularly evident in the orbito-frontal cortex and insula, with considerable expansion and age-related GM decline, while low decline and expansion was found in the basal ganglia, occipital lobe, temporal pole, and medial temporal lobe. This general association was replicated in another large dataset, the eNKI (Enhanced Nathan Kline Institute; n = 765; r = 0.51; p = 0.01)^44^ sample using the same OPNMF parcellation to extract the age-mediated GM decline (Supplementary Figure 4). Furthermore, we replicated the significant relationship between cerebral expansion and agerelated GM decline in humans applying the much finer Davi130 chimpanzee parcellation^8^ (r = -0.38, p = 1×10^-4^).

**Figure 5.**
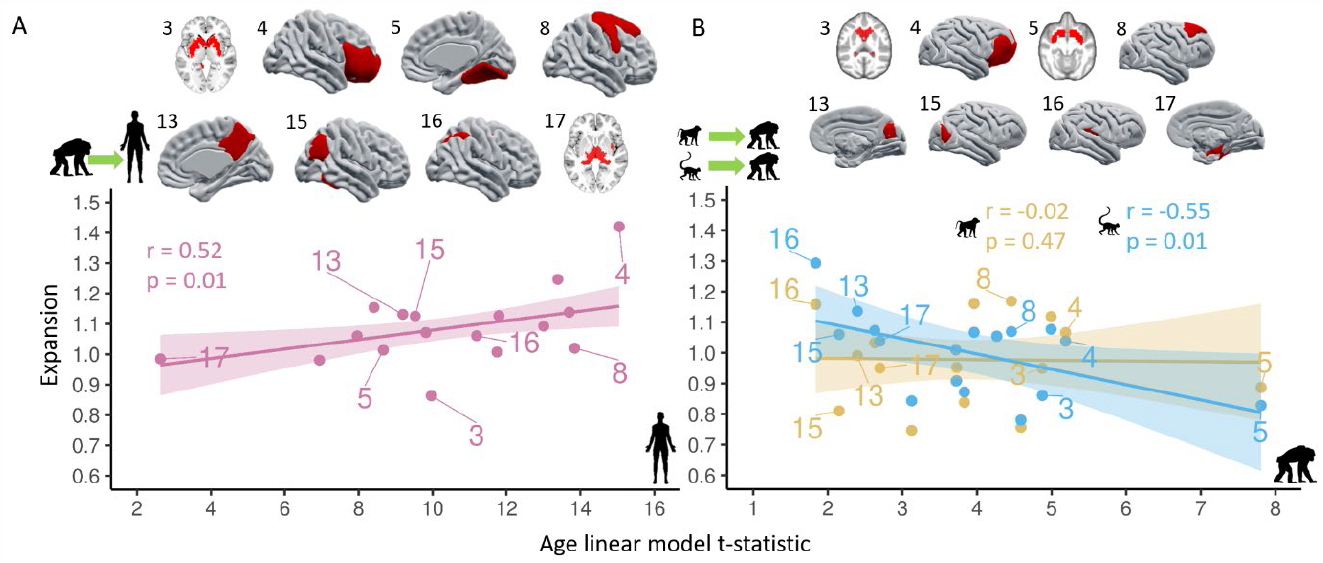
Gray Matter Aging and cerebral expansion. Each dot represents an OPNMF brain parcel and a selection of parcels for human (A) and chimpanzee (B) are shown above the scatter plots. A – Chimpanzee to human expansion and human age-related GM decline (Pink), B – macaque to chimpanzee expansion (Blue) and baboon to chimpanzee expansion (yellow) correlated with chimpanzee age-related GM decline. Significance (p) of Person’s correlation (r) for cross-species expansion and age-related GM decline relationship is determined by permutation testing (k = 100 000).

In chimpanzees, no significant relationship was seen between aging and cerebral expansion from baboon (r = -0.02, p = 0.47), but a negative correlation was found for expansion from macaque (r = -0.55, p = 0.01). Even though the macaque and baboon to chimpanzee expansion maps show a similar pattern in spatial distribution, there is an apparent trend of regions showing greater macaque to chimpanzee expansion in regions with minimal age-related GM decline. This relationship is strongly driven by the greater macaque to chimpanzee expansion in peristriate cortex, precuneus, and posterior insula as compared to baboon related expansion. Using the Davi130 parcellation^8^, both cercopithecoid monkeys to chimpanzee expansion maps showed no relationship (baboon to chimpanzee - r = 0.11, p = 0.12 and macaque to chimpanzee – r = 0.08, p = 0.21) with age-related GM decline in chimpanzees.

## Discussion

We developed a data-driven framework for interspecies comparison and found a human-specific positive relationship between age-related GM decline and expansion in the human brain compared to chimpanzees. In chimpanzees, on the other hand, there was no such brain age association for cerebral expansion relative to baboons and even a negative correlation with the brain’s expansion from macaque to chimpanzee. This suggests that the extensive expansion of the PFC and other cortical association areas in recent human evolution since splitting from a common ancestor with chimpanzees comes at the price of increased age-related vulnerability.

Unsupervised clustering of gray matter structure separately in humans and chimpanzees using OPNMF yields low dimensional spatial parcellations matching coarse macroanatomy that provides a basis for cross-species comparison of brain organization. Our approach establishes brain parcellations based on species-specific information, while identifying comparable organizational features between species. Both human and chimpanzee factorization solutions show overall hemispheric symmetry and spatial contiguity, and align with known macroanatomical structures^8^ as reported in previous applications of OPNMF in humans^28,31–33,45^. The 17-cluster solution was selected based on cross-species similarity and within-species reconstruction accuracy matching not only the granularity previously selected using OPNMF in human infants^45^ and^33^ but also resembling overall spatial structure of those OPNMF solutions. Similarities include parcels representing precuneus, insula, and superior parietal lobe, in the human factorization while both the chimpanzee and human solutions show similarities to previous findings in the PFC and temporal pole^27,30,32,44^. Additionally, the superior parietal lobe and PFC parcels show similarities to spatial clusters that were reported to represent genetic influence on cortical thickness^46^.

Independently in chimpanzees and humans, the OPNMF 17-cluster solution established a parcel representing the ventrolateral and orbital parts of the PFC, which shows the highest crossspecies similarity as well great expansion in humans relative to chimpanzees, despite the PFC being proportionally larger in humans compared to chimpanzees^11^, which is due in part to allometric scaling^47,48^. Our multivariate data-driven analysis suggests a possible similarity in the organization of the ventral and orbital sub-regions of the PFC. Furthermore, this region showed an exceptionally large age-related GM decline in humans alongside the high degree of expansion. This suggests that the greater expansion of PFC, which has been instrumental in evolutionary development in primate cognition^49^, comes with the detriment of severe age-related GM decrease in humans, where the PFC plays an important role in higher-order cognitive functions, such as executive control^24,50^ and working memory^51^. The much greater PFC expansion and age-mediated GM decline in humans compared to chimpanzees may be interpreted as additional dimension of the “last in, first out” hypothesis^26^ in the developmental maturation to aging trajectory.

The relationship between human GM volume decline and cortical expansion indicates a link between the evolutionary development of specific cortical areas in humans and increased vulnerability to neurodegenerative processes. Interestingly, such a relationship was not present in the expanded cortical regions of chimpanzee relative to baboons and macaques even though a significant GM decline was also present in chimpanzees ^8^. The main difference between humans and chimpanzees seems to be the more prominent expansion in sensorimotor regions in chimpanzees relative to the cercopithecoid monkeys, whereas regions of human cortical expansion relative to chimpanzees is generally seen in more multimodal association regions. This may be related to chimpanzees improved abilities for tool use as compared to cercopithecoid monkeys^52^. Human multimodal cortical areas are characterized by lower neuronal cell density, as well as higher dendritic branching and spine numbers of pyramidal neurons^53,54^. Interestingly, compared to other great apes, the human brain has a large neuropil fraction in the frontal pole^55^ and the anterior insula^56^. The neuropil fraction represents the space surrounding cell bodies occupied by dendritic and axonal interconnectedness through local intrinsic and extrinsic connections of a region. Both these areas (frontal pole and insula) show a combination of large expansion and age-related changes on GM volume (Fig. 4A) in humans. With dendritic reduction and synapse loss being characteristics of normal aging processes^57^, the relatively increased neuropil space of human association cortex may partly explain the aging – expansion relationship we observe. The medial and orbito-frontal cortex as well as the insula that displayed large expansion and age-mediated GM decline have been previously found to have high deterioration of glucose metabolism and large accumulation uptake of β-amyloid in human aging^58–60^.

The large expansion in frontal and parietal regions are comparable to studies using cortical surface measures to estimate the expansion from chimpanzee to human^22^ as well as macaque to human^26^. Additionally, using a latent space from functional MRI data, Xu and colleagues have also shown high expansion in parietal and frontal areas^61^, although not prefrontal that we show here. The chimpanzee to human surface expansion map showed less expansion in the ventral medial prefrontal cortex and the insula compared to our results. The difference could relate to our utilization our volumetric maps that were initially created voxel-wise and then masked with our comparative 17-cluster solution, while Wei and colleagues employed a higher granularity human surface atlas and mapped the expansion value within this atlas space. The macroanatomical surface and function expansion maps previously reported between macaque and human, both show greater expansion in the lateral temporal lobe and less in the ventral medial prefrontal cortex compared to our chimpanzee to human map. This may again relate to the different brain data type for the lateral temporal lobe, we showed greater expansion in the cercopithecoid monkey to chimpanzees, and this may show that the expansion of the lateral temporal lobe relates to monkey to great ape expansion and not specifically to humans.

Several considerations need to be considered when interpreting the results presented in this study. First, our voxel-based morphometry analysis conducted was limited to structural information present in the T1w contrast and does not include structural connectivity or functional dynamics. Such complimentary information provided by additional modalities will help to establish a more comprehensive cross-species comparison. Second, our inferences of age-related GM volume decline are driven by cross-sectional normal aging data (age ≤ 58 y/o); although a longitudinal design can provide additional information on structural change that occurs to the brain throughout the lifespan. Third, to infer the unique relationship in human evolution between aging and regions of cortical expansion we only used templates of one great ape species (chimpanzee) and two different cercopithecoid monkey species (macaque & baboon). Further research is required using additional primate species. A broader phylogenetic investigation will enable a better understanding at which evolutionary branches these aspects of the neurobiology of aging occur.

In conclusion, we propose a data-driven comparative analysis framework that reveals structural features of great ape brain organization. The species-specific GM parcellations using OPNMF contain both inter- and intra-species anatomical features and provides the basis for macroanatomical cross-species comparison. Utilizing the optimal 17-cluster solution we found a significant relationship between evolutionary brain expansion and brain aging in humans only. Regions with high cerebral expansion in humans relative to chimpanzees showed extensive GM decline. A weaker but similar pattern of brain aging in chimpanzees did not show this association with brain expansion relative to baboons nor to macaques. These findings allude to an association of recent human brain evolution, here cortical expansion to the unique vulnerability of multimodal association regions, in particular the PFC, to brain aging.

## Methods

### Sample Description

The chimpanzee T1w MRI scans were provided by the NCBR containing brain scans of 223 captive animals (137 females; 9 - 54 y/o; mean age 26.9 ± 10.2 years). The chimpanzees were housed at the National Center for Chimpanzee Care (NCC) at The University of Texas MD Cancer Center (N=147; 1.5 Tesla G.E echo-speed Horizon LX scanner) or the Emory National Primate Research Center (ENPRC; N=76; 3.0 Tesla Siemens Trio scanner). The MRI scanning procedures for chimpanzees at both the NCC and ENPRC were designed to minimize stress for the animals. Data were acquired with the approval of ethics committees at both sites and were obtained prior to the 2015 implementation of the U.S. Fish and Wildlife Service and National Institutes of Health regulations governing research with chimpanzees. Image quality control (QC) was conducted by assessing sample outliers in voxel-wise GM intensity correlations. 194 chimpanzees (130 females, 9 - 54 y/o, mean age = 26.2 ± 9.9) passed QC. To minimize the effect of extreme aging on the OPNMF solutions subjects over 50 years old were removed for a final sample of 189 chimpanzees (126 females, 9 - 50 y/o, mean age = 25.6 ± 9.1).

The human structural T1w MRI scans were provided from the IXI (Information eXtraction from Images) dataset. This open dataset was specifically chosen for comparison with the NCBR chimpanzees as it contains both 1.5 and 3 Tesla brain scans and has a comparable distribution of age and sex. IXI consists of 565 healthy subjects (314 females; 20 - 86 y/o; mean age 48.69 ± 16.46 years) without missing metadata. To further match the two NCBR scanners, subjects from the Hammersmith Hospital (HH; N=181; 3.0 Tesla Philips Medical Systems Intera scanner) and Guy’s Hospital (GUYS; N=315; 1.5 Tesla Philips Medical Systems Gyroscan Intera scanner) were considered. These 496 subjects (270 females; 20 - 86 y/o; mean age 49.57 ± 16.28 years) all passed QC. To aid in comparability to the chimpanzee sample, the very old IXI subjects (>75 y/o) were removed for the construction of the OPNMF solutions for a final sample of 480 subjects (262 females, 20 - 74 y/o, mean age = 48.7 ± 16.5). We used the eNKI open neuroimaging dataset^44^ to replicate the aging – expansion relationship in a larger lifespan sample. The eNKI scans were all acquired using a single 3T scanner (Siemens Magnetom TrioTim). T1w images were obtained using a MPRAGE sequence with 1mm isotropic voxels and TR = 1900 ms. The T1w images were preprocessed the same as IXI using CAT12. Following preprocessing, patient identified subjects were removed and QC was conducted by removing subjects that had CAT12 image quality ratings above two standard deviation from the mean. We then used 765 images from eNKI^44^ (502 females; 6 – 85 y/o; mean age = 39.8 ± 22.2) for our replication analysis.

### Image Processing

The chimpanzee (NCBR) and human (IXI, & eNKI) samples were preprocessed using the SPM12 (Statistical Parametric Mapping; https://www.fil.ion.ucl.ac.uk/spm/software/spm12/; v7487) toolbox, CAT12^62^ (Computational Anatomy Toolbox; http://www.neuro.uni-jena.de/cat/; r1725). The NCBR sample was processed utilizing the newly established chimpanzee specific pipeline^8^ while the IXI sample utilized the default human processing pipeline with high accuracy shooting regestration^63^. The general steps of preprocessing were, first, the single subject images are affine registered to template space and segmented into the three tissue types, gray matter, white matter, and cerebrospinal fluid, utilizing a tissue probability map (TPM). Next, each tissue map is nonlinearly registered to five shooting templates with increasing registration accuracy^63^ to bring all subjects’ tissue maps into the same template space. Finally, the deformation fields required to register the subjects into template space are used to modulate the tissue maps to conserve original local volume. Following preprocessing the modulated GM maps for each species were down-sampled (2 mm & 3 mm resolution) and smoothed (4 mm & 6 mm full width half maximum) in the NCBR and IXI samples, respectively. Finally, a GM mask at 0.3 and 0.2 probability for the chimpanzee and human samples respectively was applied encompassing the cortex and basal ganglia.

### Orthogonal Projective Non-negative Matrix Factorization (OPNMF) and Granularity Selection

To estimate the GM structural covariance parcellation the orthogonal variant of NMF^64^, OPNMF was used^228,29^. OPNMF establishes a low dimensional representation of the voxel-wise GM data where non-negativity is enforced on all elements. The low dimensional space comprises a parcellation matrix with the factorization loadings for each voxel and a subject matrix containing the subject loadings for each component. By constraining the matrices to non-negative values provides a parts-based representation of the cerebral GM data by the way of spatially continuous and additive covariance parcels. A hard parcellation is created by appointing each voxel to the parcel with the highest weight. Selection of the most appropriate OPNMF granularity for cross-species comparison was assessed through analysis of the reconstruction error development over bootstraps (k = 100) in combination with cross-species OPNMF solution parcel similarity over the range of 2 – 40 parcels. Further details can be found in the Supplementary.

### Template Processing and Expansion Maps

Employing CAT12 preprocessing we established cross-species deformation maps which represent an approximation of the cross-species expansion. Utilizing population average T1 maps of a species and the processing pipeline of the target species, we could create deformation maps that represent an estimation of cross-species expansion. Species templates were selected as they represent an average brain, improving interpretability of the expansion estimates as well as having high tissue contrasts aiding segmentation and registration. We used two species-specific processing pipeline, chimpanzee ^8^ and human, to create our expansion maps. The chimpanzee to human expansion map was created using the chimpanzee template in the human pipeline while the baboon and macaque to chimpanzee expansion maps were created using the chimpanzee pipeline^8^ with the baboon^36^ and macaque^37–39^ templates respectively. We averaged the expansion map of three commonly used macaque templates to encompass inter-sample variation. Further information regarding the expansion map creation can be found in the Supplementary. QC was performed on all deformation and expansion maps by assessing the smoothness and feasibility of the macroanatomical structures. Additionally, for the chimpanzee to human map we visually inspected the spatial location of the Davi130^8^ regions when registered to the human template space (Supplementary Figure 1).

### Gray Matter Aging and Expansion Analyses

We assessed the relationship between age related GM decline and relative expansion in chimpanzees and humans, utilizing the data-driven low dimensional OPNMF solution. The age-related changes on GM volume was assessed for the chimpanzee and human samples utilizing an OPNMF parcel-wise linear regression model. The age-range of the IXI sample was matched to the chimpanzee to improve comparability, by accounting for the difference in aging processes (human ≈ 1.15x chimpanzee)^41^. Therefore, 304 IXI subjects (females = 150; 20 – 58 y/o; age = 39.0 ± 11.0) and 189 chimpanzees (126 females; 9 - 50 y/o; age = 25.6 ± 9.1) were used for the age regression model. Average GM volume values for each OPNMF parcel from the chimpanzees and humans were entered into a regression model for each species, as the independent variable with age, sex, total intracranial volume (TIV), and scanner field strength as the dependent variables. Parcels showing a significant age-mediated GM decline was assessed at p≤0.05 following correction for multiple comparisons using FWE^42^. The parcel-wise age model t-statistics in the human sample were compared with the chimpanzee to human expansion while chimpanzee age-mediated GM decline was compared with the baboon and macaque cross-species expansion. Cross-species expansion was estimated by taking the mean expansion from each component for the various cross-species expansion maps and z-scored to present the inter-regional inter-species expansion. Significance of Pearson’s correlation and difference between correlations was determined through permutation testing (k=100 000) at p≤0.05. Finally, the human parcel-wise age-related GM decline was replicated in the eNKI^44^ lifespan neuroimaging dataset and correlated with chimpanzee to human expansion to assess the robustness of our findings in humans.

## Data and Code Availability

The chimpanzee (www.chimpanzeebrain.org) and two human neuroimaging datasets, IXI (http://brain-development.org/ixi-dataset/) and eNKI (http://fcon_1000.projects.nitrc.org/indi/enhanced/index.html) are openly available. The human (MNI) and chimpanzee (JunaChimp) reference templates are available as part of the CAT12 (v12.7+) toolbox or can be individually downloaded at http://www.bic.mni.mcgill.ca/ServicesAtlases/ICBM152NLin2009 and http://junachimp.inm7.de/ respectively. The cercopithecoid monkey template are also openly available for download, baboon (Haiko89 - https://www.nitrc.org/projects/haiko89/) and macaques (D99 – https://afni.nimh.nih.gov/pub/dist/doc/htmldoc/nonhuman/macaque_tempatl/atlas_d99v2.html; INIA19 – https://www.nitrc.org/projects/inia19/; NMT - https://afni.nimh.nih.gov/pub/dist/doc/htmldoc/nonhuman/macaque_tempatl/template_nmtv2.html#download-symmetric-nmt-v2-datasets). The GM masks for chimpanzees and humans can be downloaded at https://zenodo.org/record/6463123#.YyljX_exVhG. The remaining data created for this manuscript can be downloaded at https://zenodo.org/records/10141986 and code for the analyses can be found at https://github.com/viko18/GreatApe_Aging.

