## Supplementary for "The Uniqueness of Human Vulnerability to Brain Aging in Great Ape Evolution"

### Orthogonal Projective Non-negative Matrix Factorization

To extract multivariate GM factors for cross-species comparison we employed an orthogonal modification of non-negative matrix factorization (NMF), orthogonal projective NMF (OPNMF)[1,2]. NMF factorizes a data matrix (X) with dimensions m x n (here GM density voxels x No. subjects) into a factor matrix W (m x k, voxels x factors) and a subject-specific factor weight matrix H (k x n, factors x subjects), whereby all three matrices contain elements of non-negative value. The construction of the factor (W) and weight (H) matrices is achieved by minimizing the reconstruction error between the original input matrix and its reconstruction by the multiplication of the two factorized matrices (W & H). NMF establishes a parts-based representation of the original input data through the created factors[3]. Therefore, the factors represent separate interpretable parts of the multivariate input data extracted from the underlying patterns of variance.

OPNMF factorizes the data matrix by solving the minimization problem $||X - WW^{T}X||$ which is subject to $W^{T}W = I$; $W \geq 0$ where, ||.|| refers to the squared Frobenius norm and *I* denotes the identity matrix. To first initialize the W matrix for the factorization we employed non-negative double singular value decomposition (NNSVD)[4] which encourages sparsity of factors. Subsequently, *W* is iteratively updated (k=10 000) with the multiplicative update rule, $W_{ij}^{'} = W_{ij}$ until it reaches an optimal solution[2]. The final step is to project *X* onto *W* to calculate *H*.

We decided to employ OPNMF variation instead of the original NMF as it provides several advantages when representing structural T1w MRI data as a small number of structural covariance factors[1,5]. OPNMF is different from standard NMF in how it constructs the factor loading weights of the H matrix. In standard NMF, the matrix H is estimated separately while in OPNMF it is estimated by projecting the input matrix (X) onto the factor matrix (W) using $H = W^{T}X$. Therefore, in OPNMF all factors participate in the reconstruction of all data points while in NMF a subset of factors is involved in reconstructing a subset of data points leading to greatly less overlap of factors and more sparsity in OPNMF as compared to NMF. This leads to the creation of structural covariance factors that are spatially continuous with minimal overlap providing a low dimensional representation of the underlying GM data that is easier to interpret. Such features in OPNMF enable each voxel to be assigned a factor in the brain by employing a winner takes all approach when back projecting the factors back onto the brain to create the final cluster solution.

### Selected OPNMF Parcellation Solution

To determine an informative low dimensional structural covariance representation of GM tissue utilizing OPNMF we employed a data-driven approach accounting for accuracy, stability, and inter-species cluster spatial similarity. As the reconstruction error is a depiction of how well the factorization solution estimates the original input matrix, we assessed its change (decrease) as the number of factors increased to determine factor solutions that accurately represent the input data. This allows us to determine the improvement of the factorizations estimation of the original data when increasing the granularity or factor number. Therefore, a plateau in this improvement represents factor solutions where increasing the granularity has minimal improvements in the estimation of the original data. Consequently, some intrinsic and informative dimension of the data has been reached and the data modeled by further factors is more difficult to discern whether they represent signal or noise. Accordingly, we averaged the change in reconstruction error over 100 bootstrapped implementations of OPNMF across a range of factor numbers (2 – 40 steps of 1) to provide a stable indication of the accuracy change for each species separately. Additionally, to improve the inter-species comparability we selected the factor solution with the highest inter-species spatial similarity measured using adjusted rand index (ARI). The ARI was calculated following the deformation of the chimpanzee OPNMF solutions to the human MNI space. It determines the factors spatial similarity above chance with a value between zero and one. Through this approach, the 17-factor solution was selected for both chimpanzees and humans.

### Quality Control

Quality control (QC) was conducted by checking the sample inhomogeneity utilizing CAT12 (Computational Anatomy Toolbox). The modulated GM maps with a mean correlation below two standard deviations were flagged for visual inspection. The flagged images were then removed if they contained tissue misclassification, artifacts, irregular deformations, or very high intensity values. This process was repeated a second time with the passed images in the chimpanzee sample only as no images were removed in the IXI sample. Following the second iteration in the chimpanzee sample no more images were flagged. Following QC, 194 (130 females; 9–54 y/o; mean age = 26.2 ± 9.9) chimpanzees and 496 (270 females; 20 - 86 y/o; mean age 49.57 ± 16.28 years) human T1-weighted images qualified for further investigation.

To ascertain the feasibility and usability of the cross-species, chimpanzee to human, deformation maps visual QC was conducted. The chimpanzee Davi130 macroanatomical parcellation[6] was deformed to the human MNI space and visually inspected for large systematic misalignment with the expected macroanatomical structures (Supp. Fig. 1). There are some slight misalignment of gyri and the superior cerebellum and posterior part of the superior frontal gyrus have moved too far superiorly. As the OPNMF factors used in cross-species comparison are quite large and additional gray matter (GM) masking was conducted, these small differences will not greatly affect our analysis. Therefore, it shows the cross-species deformation map is able to approximate chimpanzee macroanatomical features onto a human template brain. Additionally, we visually inspected the deformed chimpanzee OPNMF factor solutions in MNI space for any large artifacts.

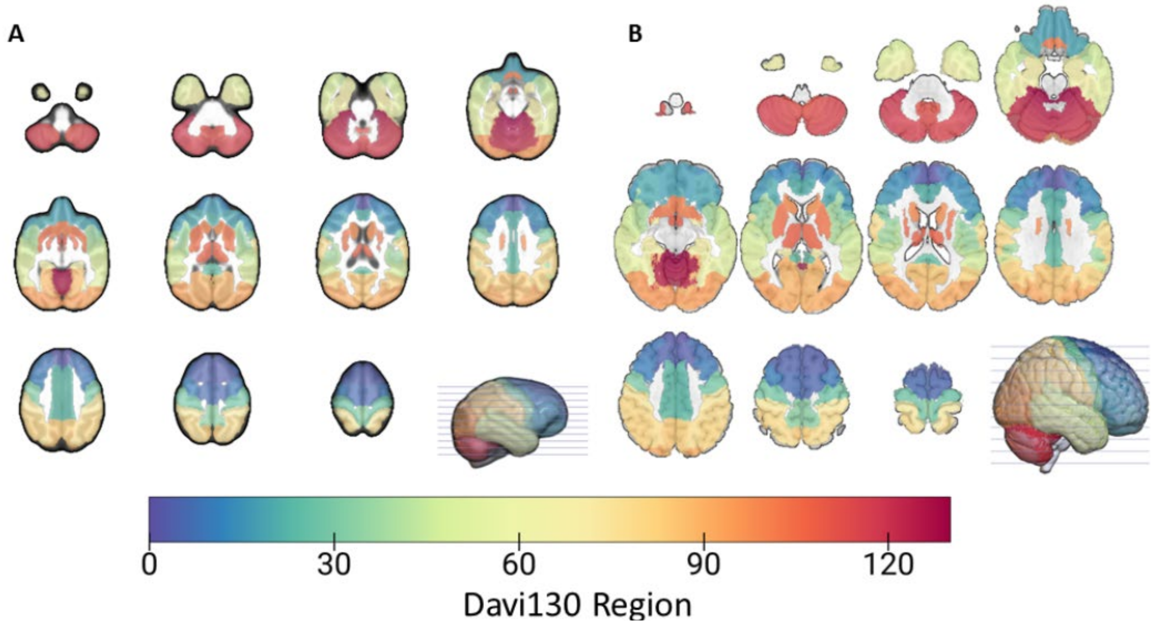


**Supplementary Figure 1. Deformation map quality control.** Davi130 chimpanzee parcellation in chimpanzee (A) and human (B) template space used for visual quality control of chimpanzee to human deformation map.

### Davi130 Aging Effect on Gray Matter

We assessed the age related changes to GM volume in chimpanzees and humans at a higher granularity than the 17-factor OPNMF solution, by using the chimpanzee macroanatomical Davi130 parcellation[6]. The chimpanzee parcellation was projected to human template space utilizing the chimpanzee to human deformation map. To be comparable with the OPNMF factor solution aging results the Davi130 cerebellum regions were not analyzed which left 110 cortical and sub-cortical regions. The same chimpanzee (n=189; 126 females; 9 – 50 y/o; mean age = 25.6 ± 9.1) and human (n=304; 150 females; 20 – 58 y/o; mean age = 39.0±11.0) samples were used as for the OPNMF factor aging analysis. The aging effect on GM volume was determined with a multiple linear regression model over all regions. Average GM for each region was used as the dependent variable with age, sex, total intracranial volume (TIV), and scanner field strength as independent variables. Significant age effect was determined at $p \leq 0.05$ following correction for multiple comparisons using family wise error (FWE)[7]. The Davi130 aging effect (Supp. Fig. 2) shows a similar spatial pattern as the OPNMF 17-factor solution results in both species. In the chimpanzee, significant aging effect was found in the basal ganglia, cingulate cortex, lateral temporal lobe, frontal cortex, and precuneus. This is considerably less than the OPNMF result due to the higher threshold for significance following multiple comparison correction with the higher amount of regions. The largest aging effect was found in the caudate nucleus and the lowest in the motor cortex and occipital lobe comparable to the OPNMF aging result. In the human sample the Davi130 aging result showed similar large aging effect in the frontal and lateral temporal cortex and low effect in the basal ganglia, in particular the thalamus.

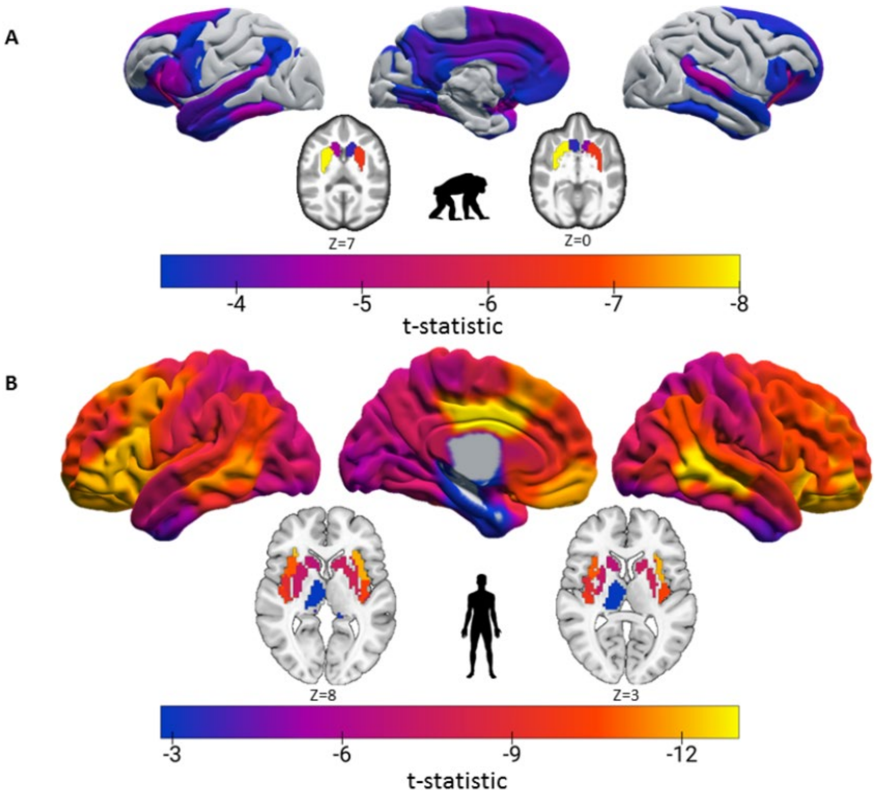


**Supplementary Figure 2. Comparison of aging effect on GM volume.** Aging effect on GM volume across all cortical and sub-cortical Davi130 regions in chimpanzees (A) and human (B) samples. Significant regions at p≤0.05 are presented following correcting for multiple comparisons using FWE.

### Expansion map creation

We imported a species average T1 map (e.g. chimpanzee – JunaChimp[6]) into the pipeline for the other species (e.g. human – MNI[8]) to create maps of the non-linear registration across species. Following processing the JunaChimp[6] chimpanzee template using the standard the CAT12 human pipeline the deformation field map was used to register the chimpanzee OPNMF parcellations to MNI space to conduct similarity analysis for granularity selection. To create the relative expansion maps, we used the modulated Jacobians from processing and conducted post-hoc manipulation by masking (brain mask) and converting the Jacobian values into cross-species expansion approximations. The expansion maps were created by dividing the jacobians by a scaling factor, which was the inverse of the relative difference in brain size between species. Therefore, as the human brain is approximately 3.5x larger than the chimpanzee a scaling factor of 0.286 (1/3.5) was applied. The chimpanzee brain is approximately 2.5x larger than the baboon and 4.5x the macaque so a factor of 0.4 and 0.22 were applied respectively.

### Brain Aging and Cross-species Expansion Comparison Replication

**Davi130 Parcellation**

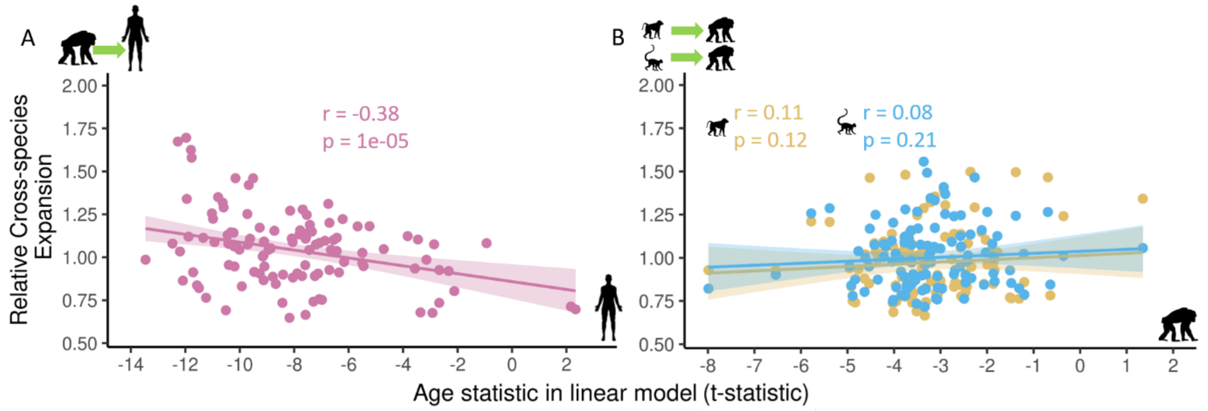


**Supplementary Figure 3. Aging – expansion comparison (Davi130).** Scatter plots showing of cross-species expansion and aging effect between A – chimpanzee to human expansion and human aging effect (Pink), B – macaque to chimpanzee expansion and chimpanzee aging effect (Blue) and baboon to chimpanzee expansion and chimpanzee aging effect (Yellow). Significance (p) of correlation (Person's r) for cross-species expansion and aging effect relationship is determined by permutation testing (k = 100 000).

**eNKI Human Lifespan Sample**

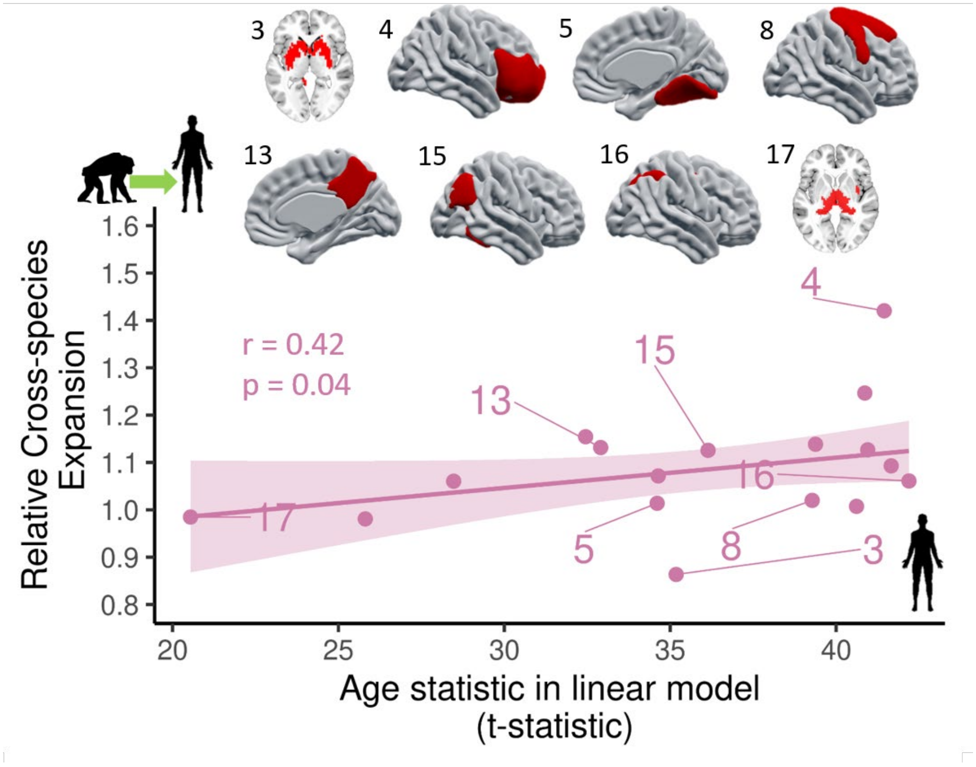


**Supplementary Figure 4. Aging – expansion comparison (eNKI).** Scatter plots showing of cross-species expansion and aging effect using the IXI sample OPNMF 17-factor solution to extract aging effect for each factor using the eNKI sample (n=765, 502 females; 6 – 85 y/o; mean age = 39.8 ± 22.2). A selection of OPNMF factors are projected onto volume slice or rendering of the MNI human template.

Significance (p) of correlation (Person's r) for cross-species expansion and aging effect relationship is determined by permutation testing (k = 100 000).
